# Phylogenetic distribution and experimental characterization of corrinoid production and dependence in soil bacterial isolates

**DOI:** 10.1101/2023.12.21.572947

**Authors:** Zoila I. Alvarez-Aponte, Alekhya M. Govindaraju, Zachary F. Hallberg, Alexa M. Nicolas, Myka A. Green, Kenny C. Mok, Citlali Fonseca-Garcia, Devin Coleman-Derr, Eoin L. Brodie, Hans K. Carlson, Michiko E. Taga

## Abstract

Soil microbial communities impact carbon sequestration and release, biogeochemical cycling, and agricultural yields. These global effects rely on metabolic interactions that modulate community composition and function. However, the physicochemical and taxonomic complexity of soil and the scarcity of available isolates for phenotypic testing are significant barriers to studying soil microbial interactions. Corrinoids—the vitamin B_12_ family of cofactors—are critical for microbial metabolism, yet they are synthesized by only a subset of microbiome members. Here, we evaluated corrinoid production and dependence in soil bacteria as a model to investigate the ecological roles of microbes involved in metabolic interactions. We isolated and characterized a taxonomically diverse collection of 161 soil bacteria from a single study site. Most corrinoid-dependent bacteria in the collection prefer B_12_ over other corrinoids, while all tested producers synthesize B_12_, indicating metabolic compatibility between producers and dependents in the collection. Furthermore, a subset of producers release B_12_ at levels sufficient to support dependent isolates in laboratory culture at estimated ratios of up to 1,000 dependents per producer. Within our isolate collection, we did not find strong phylogenetic patterns in corrinoid production or dependence. Upon investigating trends in the phylogenetic dispersion of corrinoid metabolism categories across sequenced bacteria from various environments, we found that these traits are conserved in 47 out of 85 genera. Together, these phenotypic and genomic results provide evidence for corrinoid-based metabolic interactions among bacteria and provide a framework for the study of nutrient-sharing ecological interactions in microbial communities.

## INTRODUCTION

Microbes engage in metabolic interactions that collectively define ecological networks in communities (1). Microbial interactions can be key mediators of community function, and disruptions to interactions can restructure whole communities (2,3). Thus, it is crucial to disentangle microbial interactions and generate a predictive understanding of nutritional influences on communities and, in turn, on the environment.

Experimental and computational studies have shown that microbes commonly lack the ability to synthesize all of the metabolites they require (4,5). For example, many microbes are unable to synthesize certain cofactors and amino acids, and therefore must acquire these nutrients from other organisms in the environment (6–8). As a consequence, microbial communities are composed of “producer” and “dependent” organisms that synthesize and require a given nutrient, respectively. The complexity of microbial communities is due, in part, to this network of interactions arising from interdependence among microbes that produce and require a range of different nutrients. Such interdependence may develop because loss of biosynthesis genes can be evolutionarily favored in contexts where required nutrients are abundant in the environment or can be acquired from other microbes (9,10).

Investigating the molecular mechanisms and community impacts of nutritional interactions is challenging for many reasons. First, many microbial communities are functionally diverse and contain numerous metabolites that are produced, used, and chemically transformed by community members. Second, while some metabolic capabilities can be inferred from genomic data, these analyses currently lack spatial and temporal resolution, making it difficult to predict how interactions among community members may be impacted by the metabolic activities of a single microbe. Further, because most microbes have not been isolated in pure culture (11), metabolic predictions of the uncultured majority have yet to be confirmed. The challenges common to microbiome studies are amplified in soil due to its taxonomic diversity, physical and chemical heterogeneity, and environmental fluctuations, as well as disturbances due to animal or human activities (12–14). Nonetheless, generating mechanistic knowledge of nutrient cycling in soil communities is essential because of their broad impacts on the health of our planet (15,16). We address these challenges by studying the ecological roles of microbes in relation to one class of model shared nutrients in a collection of newly isolated soil bacteria.

We took a reductionist approach to investigate nutrient production and dependence by focusing on corrinoids as a representative class of shared metabolites. Corrinoids are produced by a subset of bacteria and archaea and include the vitamin B_12_ (cobalamin) family of cobalt-containing cobamide cofactors and their biosynthetic precursors (Fig. 1A). Corrinoids are required cofactors for methionine synthesis and propionate metabolism in most eukaryotes, and additionally are used by prokaryotes for other diverse processes such as mercury methylation, natural product biosynthesis, nucleotide synthesis, and numerous carbon and nitrogen transformations (17). Corrinoid-sharing interactions between producer and dependent microbes have been observed in laboratory co-cultures of bacteria (18–20), bacteria-microeukaryote pairs (21–23), and in higher-order consortia (24), and are thought to be prevalent in the gut microbiome (7).

**Figure 1.**
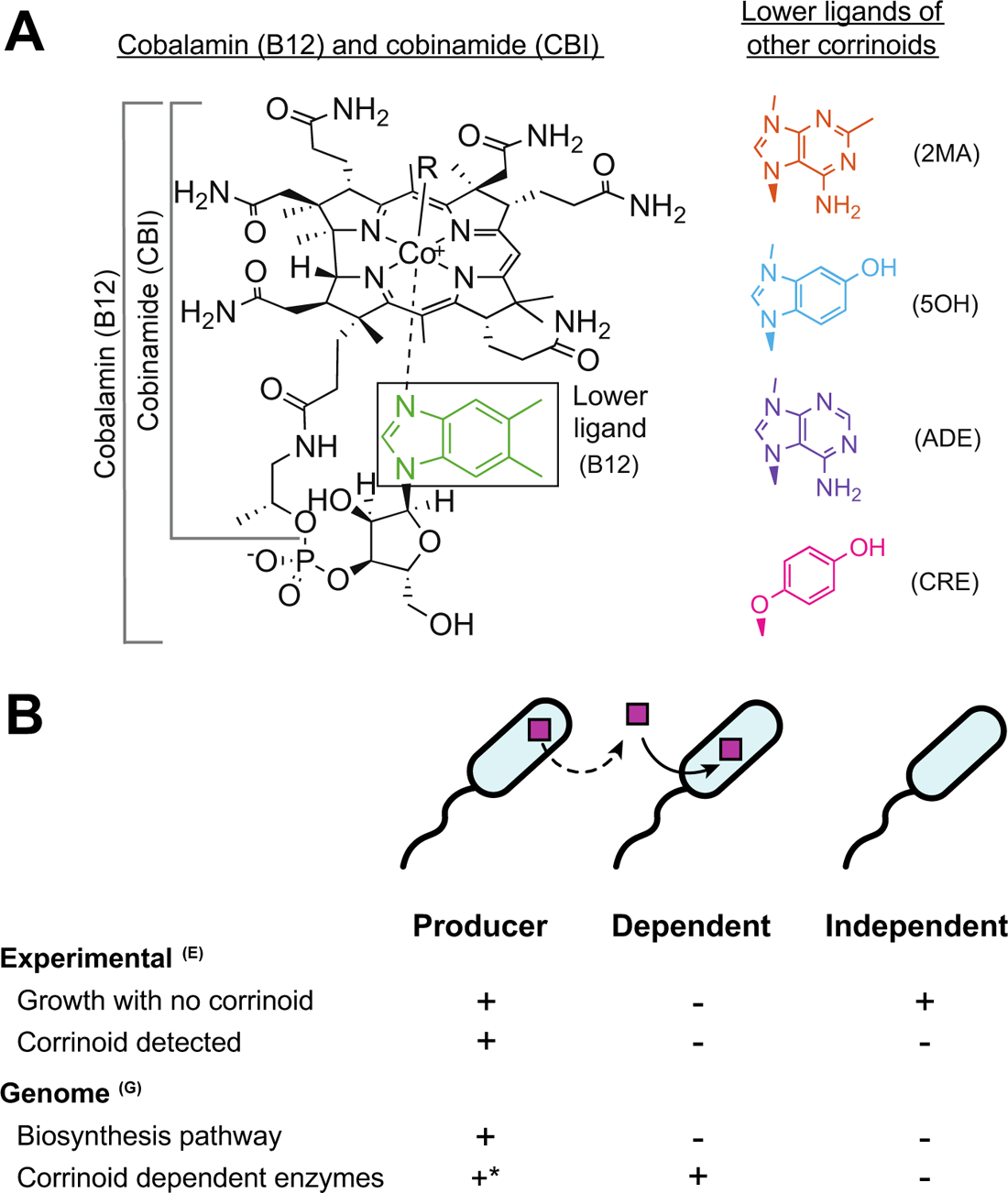
Diversity of corrinoid lower ligands and metabolic roles. (A) Chemical structure of cobalamin (B12), lower ligands of other corrinoids used in this research, and 3-letter abbreviations. From the top, 2-methyladeninylcobamide (2MA), 5-hydroxybenzimidazolylcobamide (5OH), adeninylcobamide (ADE), and cresolylcobamide (CRE). Cobinamide (CBI), shown on the left, is an incomplete corrinoid that does not contain a lower ligand. (B) Corrinoid metabolism categories include producers, dependents, and independents. Producers may release corrinoids (dashed line). These categories can be assigned based on experimental results, denoted with superscript E, and genomic analysis, denoted with superscript G, as summarized in the table. *In principle, producers^G^ are defined based solely on the presence and completeness of the biosynthesis pathway, but all corrinoid producer^G^ genomes examined thus far encode one or more corrinoid-dependent enzymes (25).

**Figure 2.**
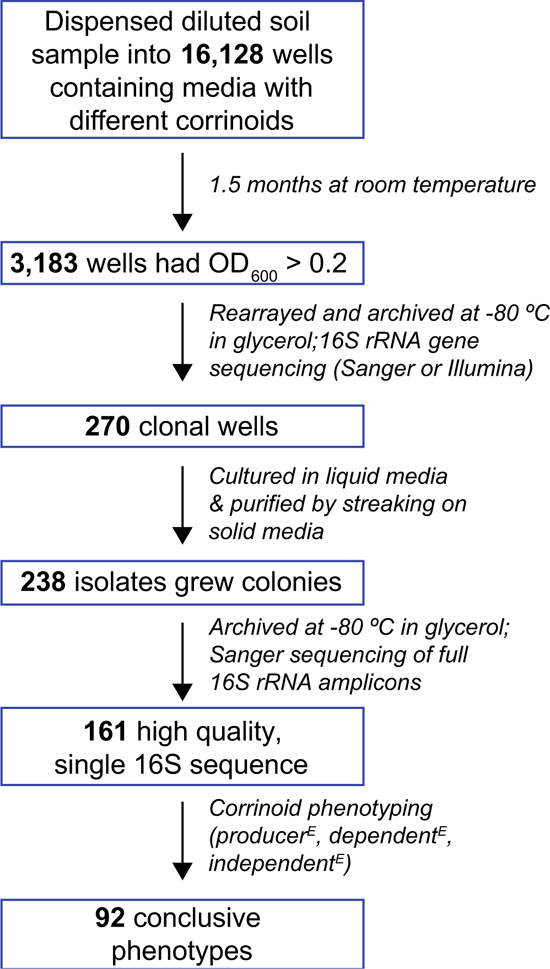
Overview of the experimental procedure. Soil bacteria were isolated by the limiting dilution method on media containing different corrinoids. Growth was detected based on OD_600_ and clonal isolates were distinguished from mixed cultures by 16S rRNA amplicon sequencing. After several purification steps, the final isolate collection contains 161 clonal isolates, of which 92 were successfully characterized as corrinoid producer^E^, dependent^E^, or independent^E^.

Computational predictions further support the hypothesis that corrinoids are broadly shared, as 37% of sequenced bacterial species are predicted corrinoid producers and 49% are dependents (25). The remaining 14% are predicted not to produce or use corrinoids, fulfilling their metabolic needs via corrinoid-independent pathways, and are therefore considered “independents” (Fig. 1B) (25). Given that dependents and producers coexist in the same environments (26–28), we hypothesize that many sharing interactions have yet to be described.

An aspect of corrinoids that influences their function as shared nutrients is their structural diversity (Fig. 1A), which has been shown to impact function – microbial preferences for particular corrinoid structures are apparent in their differential growth responses to corrinoids (17,29). Distinct groups of corrinoids have been detected in a variety of host-associated and environmental microbial communities, suggesting that microbes encounter diverse corrinoids in nature (30). Because most corrinoids are not commercially available, nearly all research on corrinoids has been performed only with B_12_ (17). Given that some microbes require corrinoids other than B_12_ (_31_), it is likely that novel bacteria that could not have been isolated on B_12_ will be culturable on other corrinoids.

In this study, we have extracted and purified four non-commercially available corrinoids to investigate the impact of corrinoid structure on bacterial growth and isolation from a California annual grassland soil. Previous work showed that this soil community contains an abundance of predicted dependents, with producers and independents in the minority (26). Consistent with these computational predictions, we found producers, dependents, and independents among our diverse collection of 161 bacterial isolates. We did not observe strong phylogenetic trends in corrinoid production and dependence in this collection, yet through our genomic analysis across the bacterial domain, we found conservation of these traits in over half of the bacterial genera we examined. Upon characterizing the corrinoid requirements of dependent isolates and corrinoid biosynthesis by producers, we found that the corrinoid synthesized by producers is compatible with the corrinoid requirements of all dependents in the collection. Further, corrinoids are released from cells in a subset of producers at levels that exceed the requirements of dependents in culture by up to 1,000-fold. These results provide an ecological framework for understanding nutritional interactions in soil through the lens of corrinoids.

## METHODS

### Isolation of bacteria from soil by the limiting dilution method

We collected soil samples from top 10 cm of an annual grassland at the Hopland Research and Extension Center in Hopland, CA (39.004056 N 123.085861 W) in November 2019 at the start of the plant growing season and April 2021 when plants typically approach peak productivity. Site characteristics and soil physicochemical properties for our sampling site were previously documented (32,33). Soil pH, determined by preparing a slurry with 1 part soil to 2 parts deionized water and measuring (n=3) with an Orion Star A111 pH Meter (Thermo Scientific, Waltham, MA, USA), was 5.87 ± 0.42 in November 2020 and 6.28 ± 0.07 in April 2021.

To separate microbial biomass from the soil, we resuspended 2.5 g of soil in 25 ml phosphate buffered saline with 2.24 mM sodium pyrophosphate (Alfa Aesar, Heysham, England), stirred for 30 minutes, and allowed the slurry to settle for 15 minutes before diluting the supernatant. All isolations and subsequent growth steps were performed using a modified VL60 medium (pH 6.0), (34) and Supplemental Table 1) amended with 0.1 g/L each of xylose, xylan, N-acetylglucosamine, and glucose, as well as 10 nM corrinoid when indicated. Cultures were grown at room temperature unless otherwise noted. The corrinoids 2MA, 5OH, and ADE were produced by guided biosynthesis in *Propionibacterium acidi-propionici* and CRE in *Sporomusa ovata*; extracted from bacterial cultures; and purified as previously described (35,36). We conducted a most probable number (MPN) count to determine the soil slurry dilution required to reach growth in approximately 30% of wells, a density that is expected to yield 80% clonal cultures, based on a Poisson distribution and previously reported isolations from human stool samples (37). 80 µl aliquots of the soil solution diluted in each medium were dispensed into the wells of 384-well plates using a Biomek liquid handler (Beckman Coulter, Indianapolis, IN, USA). Three plates per corrinoid condition were inoculated, and an uninoculated plate was prepared for each condition, for a total of 28 plates. Plates were covered with BreatheEasy (Diversified Biotech, Dedham, MA, USA) membranes for this and all subsequent steps, and incubated statically at room temperature for 44 days. Despite using the MPN calculation to determine the dilution, a surprisingly low number of wells showed growth in the November 2019 experiment (2.4%). Cultures from wells in which the OD_600_ (measured on a Tecan Spark plate reader (Gr<u>ödig</u>, Austria)) exceeded 0.29 were transferred into fresh medium at the end of the initial incubation and grown for up to 40 days. The cultures were then split into two portions, one stored at −80°C in 25% glycerol and another prepared for sequencing. The full 16S rRNA gene was amplified by PCR from each well with primers 27F and 1492R (38) (IDT, Coralville, Iowa, USA) and DreamTaq polymerase (Thermo Scientific). PCR purification and Sanger sequencing of all amplicons using the same primers was done at the UC Berkeley DNA Sequencing Facility. Sanger sequence trimming with a 0.01 error probability cutoff and *de novo* assembly of reads were performed on Geneious Prime (2022.1.1). Cultures with a single, high-quality 16S sequence were considered clonal.

Prior to the second isolation from soil collected in April 2021, the soil sample was stored at 4°C for one month, brought to 20% moisture from an original 3.33 ± 0.58% with sterile deionized water, and incubated for one week. The plates were incubated at room temperature for 49 days. Because the percentage of wells showing growth was much higher than in the previous isolation round (37% of total wells), Illumina sequencing of the 16S V4/V5 region, amplified with the 515F and 926R primers (39,40), was performed with in-line dual Illumina indices (41,42) to identify cultures containing a single 16S sequence. The amplicons were sequenced on an Illumina MiSeq with 600 bp v3 reagents. Reads were processed with custom Perl scripts implementing Pear for read merging (43), USearch (44) for filtering reads with more than one expected error, and demultiplexing using inline indexes and Unoise (45) for filtering rare reads and chimeras. 16S sequences were searched against the RDP database (46) to assign taxonomy.

Liquid cultures prepared from glycerol stocks were purified by streaking on 2X modified VL60 solidified with 14 g/L Difco noble agar (BD, Sparks, MD, USA). Nystatin (63 ng/ml) was added to the medium in cases where fungal growth was observed. Cultures were serially purified by streaking until all observed colonies were of uniform morphology. For each isolate, liquid cultures inoculated from a single colony were stored at −80 °C in 40% glycerol. After purifying, the identity of each isolate was confirmed by a second round of Sanger sequencing.

We identified 23 sequences with higher than 99% pairwise identity to a sequence in an uninoculated well. These were considered potential contaminants and removed from the dataset. After removal of isolates with chimeric sequences, the final collection is composed of 161 isolates.

Growth curves were generated to classify isolates into groups based on the time they required to reach saturating growth (24, 48, 168, or 336 hours). Isolates were inoculated from glycerol stocks into 96-well plates in triplicate and grown for 168 hours at 28 °C, shaking at 800 rpm in a plate shaker (Southwest Science, Roebling, NJ, USA), and separately at room temperature with no shaking. Growth curves were generated by measuring OD_600_ at 0, 6, 12, 24, 36, and 48 hours, and every 24 hours until 168 hours for the shaken cultures and 216 hours for standing cultures. Because isolates grew more consistently in the shaking condition, cultures were shaken at 28 °C for all subsequent steps.

### Experimental characterization of isolates as corrinoid producers, dependents, or independents

To determine whether isolates were dependent on corrinoids for growth, isolates in the 24-, 48-, and 168-hour groups were inoculated into 96-well plates with 200 µl of media containing the corrinoid used for isolation. Following growth to saturation, each culture was diluted into two wells, one amended with the same corrinoid and the other with no corrinoid, using a multi-blot replicator that transferred approximately 3 µl per well (V&P Scientific, San Diego, CA, USA). Cultures were serially passaged three additional times into the same media to eliminate corrinoid carryover. OD_600_ was measured before and after each passage. Isolates that did not grow reproducibly in media with corrinoid were not pursued further (24 isolates). Isolates that continued to grow in media with corrinoid but stopped growing after being transferred into media with no corrinoid were classified as dependents^E^, while those that continued to grow in both conditions were considered to be either producers^E^ or independents^E^ (Fig. 1B). To evaluate the effect of corrinoids on the growth of isolates, we calculated the corrinoid-specific growth enhancement as log_2_ [1+((OD _with corrinoid_ – OD _no corrinoid_)/(OD _no corrinoid_))] (47) and determined a threshold for corrinoid dependence based on the growth of bacteria isolated in the no corrinoid condition that also underwent serial transfer (maximum value obtained from the equation plus standard deviation). If two or three of the three replicates were classified as corrinoid dependent, corrinoid dose-response assays were performed as described in ref. (48) to confirm dependence and determine the corrinoid preferences of each isolate. Curve fits for dose-response curves were performed using a four-parameter non-linear fit on GraphPad Prism (v9.5.1).

To distinguish producers^E^ from independents^E^ (Fig. 1B), 100 µl of each culture were collected at the end of the fourth passage with no corrinoid addition and lysed by incubating at 98 °C for 20 minutes. An *E. coli*-based corrinoid detection bioassay was conducted as previously described (35) to determine the presence or absence of corrinoid in each sample. Data were processed to yield a ‘growth due to corrinoid’ metric by subtracting growth due to methionine (as measured by the Δ*metE*Δ*metH* control strain) from growth of the Δ*metE* bioassay strain and normalizing to growth of the wildtype *E. coli* strain. An isolate was characterized as a producer^E^ if the normalized result was greater than or equal to 2 or if the OD_600_ of the Δ*metE* bioassay strain was greater than or equal to 0.1. Conversely, an isolate was characterized as a non-producer, and thus an independent^E^, if the normalized result was less than 2 and the *E. coli* Δ*metE* OD_600_ was less than 0.1. Our method was validated using a set of previously isolated soil bacteria (49) that were genomically predicted to be corrinoid producers (Supplemental Table 2). Isolates that repressed growth of *E. coli* (2 isolates), grew to an OD_600_ less than 0.1 (7 isolates), or for which the three replicates or results for the dependence and production were inconsistent (11 isolates) were deemed inconclusive.

Data processing and analysis were performed using Python 3.7 on Jupyter Notebooks (version 6.2.0).

For further characterization of producers, we grew 1 L cultures of each in VL60 medium with no corrinoid and 200X amino acids and extracted corrinoids as described previously (35). Corrinoid extracts were analyzed by high-pressure liquid chromatography (HPLC) on a 1200 series HPLC system equipped with a diode array detector (Agilent Technologies, CA, USA) and compared to authentic corrinoid standards using Method 2, as described previously (31).

### Genus-based predictions of corrinoid metabolism

We used the dataset developed by Shelton et al. (25) which reports corrinoid biosynthesis and dependence predictions for 11,436 bacterial species. Species that were previously classified as very likely, likely, or possible producers were considered producers (25). Corrinoid-dependent species were defined as those previously classified as very likely or likely non-producers that had at least one corrinoid-dependent function, regardless of whether their genomes encoded specific corrinoid-independent alternative enzymes. Corrinoid-independent species were defined as those that were likely or very likely non-producers and had no corrinoid-dependent functions. After defining these categories, we grouped the species into their respective genera (JGI IMG taxonomic metadata was downloaded on July 18, 2023, to update any reclassified genomes). To establish a reliable cutoff for our predictions, we chose genera containing 20 species or more and made a prediction when 95% or more of the species in a genus fell under the same category.

### Phylogenetic tree building

The phylogenetic tree of the isolates in Figure 3 was constructed from full-length 16S sequences. Taxonomy assignment and tree building were done using the Silva Alignment, Classification, and Tree (ACT) service (50). To determine whether isolates were likely novel, we used BLAST to search assembled sequences against the NCBI Reference Database using Geneious (2022.1.1). If the pairwise identity between the isolate sequence and the top hit was lower than 98.6%, we considered the isolate to be novel (51).

**Figure 3.**
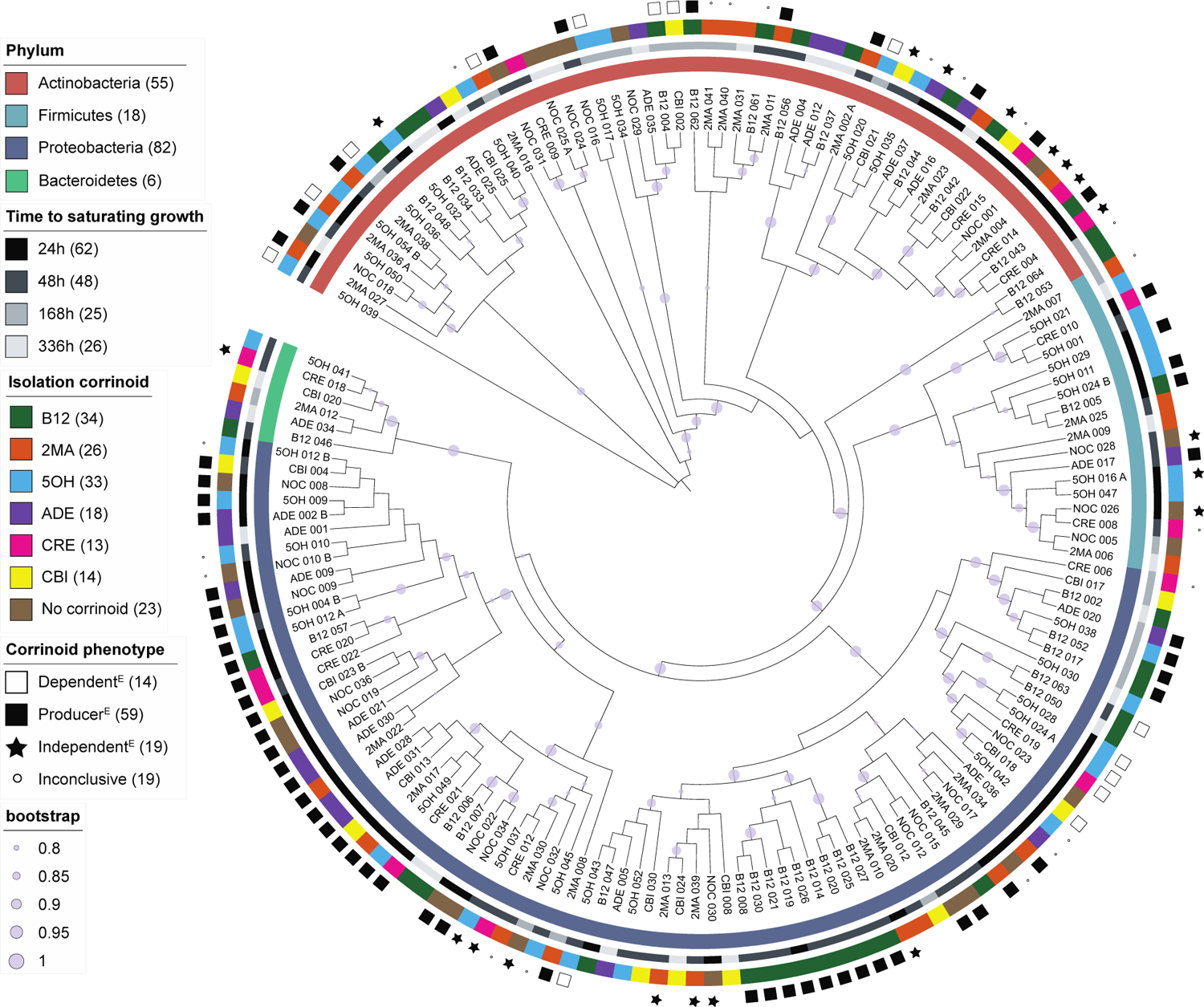
Overview of the isolate collection. A phylogenetic tree was built from the full-length 16S rRNA gene sequences of the isolates. The six- to seven-character ID is shown for each isolate. Light purple circles show bootstrap values above 0.8. Rings from the inside out show, for each isolate, (1) phylum as identified by SILVA taxonomy, (2) time to saturating growth determined from growth curves, (3) the corrinoid used in the isolation medium, and (4) the experimentally determined corrinoid metabolism category. Numbers in parentheses correspond to the number of isolates belonging to each category.

The phylogenetic tree in Figure 4 was generated using the full-length 16S sequences for the type species of each genus (85 species) and 35 additional type species that were added for context and later pruned. Sequences obtained from the NCBI Reference Sequence database can be found in Supplemental Table 3, and the phylogenetic tree including all species and bootstrap values can be found in Figure S3. A MUSCLE (52) alignment and FastTree (53) were used to generate the tree on Geneious (2022.1.1) using default settings. Tree pruning and annotation for both trees were performed on iTOL (54).

**Figure 4.**
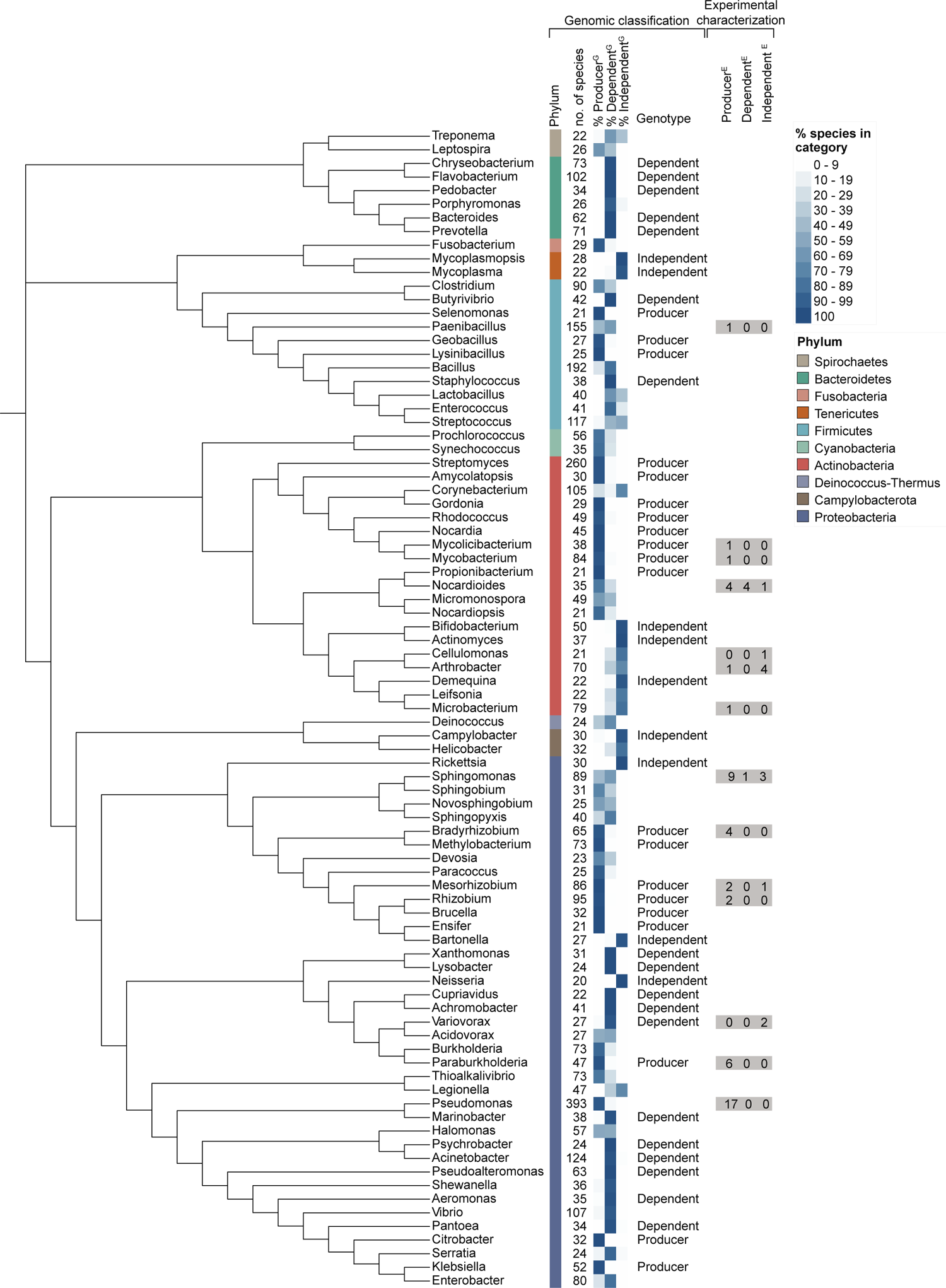
Genome-based predictions of corrinoid metabolism at the genus level. The phylogenetic tree was built from the full-length 16S sequences of the type species of 85 genera from the dataset in Reference (25) that met our cutoff by having 20 species or more. The first two columns show the phylum and the number of species analyzed for each genus, respectively, which total 10 phyla and 4,720 species. The next three columns show the percent of species in each genus predicted to belong to each corrinoid metabolism category. A corrinoid-specific genotype is indicated if 95% or more species in a genus belong to the same category. The columns labeled Experimental characterization show the number of isolates in the collection found to belong to each category based on experimental results. The unpruned tree that was used to generate the figure is shown in Fig. S3.

## RESULTS

### Generating a collection of 161 bacterial isolates from soil

To generate a collection of soil bacterial isolates with a diversity of corrinoid requirements, we performed the limiting dilution method (37) in 384-well plates containing media with one of six different corrinoids (Fig. 1A) or no corrinoid. A first set of 8,064 wells yielded only 2.4% of wells with detectable growth; a second set of 8,064 wells yielded 37%, totaling 3,183 wells with microbial growth. 16S sequencing of the resulting cultures revealed that, although we expected approximately 80% of cultures to be clonal by statistical metrics alone, only 47% and 5.8% of cultures in the two sets, respectively, were clonal. This suggests that cell aggregation is more prevalent in soil than in the gut environment, where the same method led to the statistically predicted result (37). 20 phyla were found in the total collection (Fig. S1), but among clonal wells six phyla were present, of which 238 isolates representing four phyla could be revived from frozen stocks. After purifying and archiving the clonal cultures, our collection contained 161 bacterial isolates which were used for subsequent analyses (Fig. 2).

The isolate collection is dominated by the phyla Proteobacteria and Actinobacteria, with fewer representatives from the Firmicutes and Bacteroidetes phyla (Fig. 3). This is similar to the relative abundances observed in bulk soil, where Proteobacteria and Actinobacteria are the dominant phyla (33,55,56). Of the 161 isolates, 23% (37 isolates) were considered to be novel species (51). The collection comprises 31 genera and 121 unique 16S sequences, with 11 genera each represented by a single isolate and three genera represented by 18 or more isolates. Despite this diversity, we have not sampled the bacterial diversity in this soil exhaustively (Fig. S2).

### Taxonomic and phenotypic characterization of the isolate collection

To investigate whether there were phylogenetic trends for the observed phenotypes, we constructed a phylogenetic tree of the isolate collection annotated with the characteristics of each isolate (Fig. 3, Supplemental Table 4). We did not observe strong phylogenetic trends in the time required for each isolate to reach saturating growth, except that some clades of Proteobacteria contained only fast-growing isolates. Similarly, we did not observe a correlation between phylogeny and the corrinoid used for isolation. An exception was a clade of producers within Proteobacteria, all *Sphingomonas*, that were isolated on B12. Interestingly, the number of isolates recovered in B12, 5OH, and 2MA was higher than the number of isolates in the no corrinoid (NOC) condition, while ADE, CBI, and CRE led to the recovery of fewer isolates than NOC (Fig. 3).

### Classifying corrinoid metabolism phenotypes in the isolate collection

Next, we experimentally classified each isolate as a corrinoid producer, dependent, or independent. We first assessed growth in the presence and absence of corrinoid. Isolates that stopped growing following serial transfer into media without corrinoid were classified as dependents^E^, where the superscript E refers to experimental results, in contrast to genomic predictions (superscript G, discussed below). Isolates that could grow in the absence of corrinoid were tested for corrinoid production using an *E. coli-*based corrinoid detection bioassay (35) to distinguish producers^E^ from independents^E^ (Fig. 1B; see Materials and Methods). Based on these results, all three categories are represented in the isolate collection, with the majority (64%) classified as producers^E^ (Fig. 3). The abundance of producers^E^ in our collection contrasts with genome-based predictions that dependents outnumber producers across bacteria and specifically in soil (25,26,57).

To investigate potential phylogenetic trends in corrinoid metabolism categories, we overlaid the experimentally determined corrinoid phenotypes onto the phylogenetic tree (Fig. 3). At the phylum level, we observed mixed phenotypes. For example, all three categories are represented among characterized Actinobacteria and are interspersed across several clades. In contrast, in the Proteobacteria the corrinoid phenotypes are largely consistent with phylogeny. Most Proteobacteria clades are composed of only producer isolates, aside from one clade containing genera *Phenylobacterium* and *Caulobacter* that is composed of only dependents, while in a few other clades the phenotypes are interspersed. Thus, while trends are seen in some closely related isolates, large-scale phylogenetic trends in corrinoid phenotype are not apparent.

### Corrinoid metabolism is conserved in a subset of genera, enabling taxonomy-based metabolic predictions

After investigating phylogenetic trends across isolates, we explored the extent to which phylogenetic trends exist across bacteria. We analyzed our previously published corrinoid metabolism classifications for over 11,000 bacterial species (25) to distinguish between the competing hypotheses that 1) corrinoid production, dependence, and independence show strong phylogenetic trends, enabling predictions of corrinoid metabolism based on taxonomy, or 2) corrinoid metabolism categories are phylogenetically interspersed, making it impossible to infer corrinoid-related ecological roles based solely on taxonomy. In our previous study, trends were not apparent at the phylum level except in the Bacteroidetes, which were nearly all dependents^G^ (25). Therefore, we aimed to evaluate trends at lower taxonomic levels, starting with the genus level.

We searched for phylogenetic trends among the 85 genera in our dataset and classified each genus as producer^G^, dependent^G^ or independent^G^ when possible. A corrinoid metabolism category could be assigned with high confidence for 47 out of 85 genera (Fig. 4, Supplemental Table 5). In the remaining 38 genera, a single corrinoid metabolism category does not predominate, so corrinoid metabolism classifications could not be made.

To evaluate trends across higher taxonomic levels, we mapped the genomic predictions onto a phylogenetic tree constructed from the full 16S sequences of the type species for each genus (Fig. 4, Fig. S3). As expected, five of the six Bacteroidetes genera were predicted to be dependent^G^ and only one Bacteroidetes genus has a small percentage of independent^G^ species. We observed phylogenetic trends at levels higher than the genus level in some cases. The Actinobacteria form two distinct clades, one dominated by producers^G^ and the other by independents^G^. Interestingly, no Actinobacteria genera were classified as dependents^G^, although some genera have low percentages of dependent species. All genera in one Proteobacteria clade are classified as producers^G^, with the notable exception of *Bartonella*, which has undergone genome reduction (58) and is classified as independent^G^. However, other Proteobacteria clades were mixed, aside from a few sister taxa that share corrinoid genotypes in some instances, such as *Rhizobium* and *Brucella* which are both producers^G^ or *Xanthomonas* and *Lysobacter*, which are both dependents^G^. For phyla that had fewer genera, the few classified genera were independent^G^. This may be due to a bias in the dataset, which is composed of over 90% cultured bacteria that may be less likely to be dependent. Future analysis of metagenomes may reveal more dependence among phyla with fewer sequenced representatives.

Upon comparing the genomic classifications to our experimental results for the isolate collection, we found that 19 isolates belong to genera for which genomic classifications were possible. All isolates except one matched the genomic classification (Fig. 4). The exception was a *Mesorhizobium* isolate that was predicted to be a producer^G^ but found to be independent^E^, suggesting either that it is not capable of synthesizing a corrinoid or did not produce a detectable amount under our growth conditions. The confirmation of our genomic classifications with experimental data from our isolate collection lends support to the phylogenetic predictions made for certain genera.

### Corrinoid preferences of dependent isolates reveal diverse corrinoid use capabilities

Our observation that the corrinoid used for isolation did not correlate with phylogeny (Fig. 3) led us to investigate the corrinoid preferences of the 14 dependent isolates in our collection. We measured growth in media containing a range of concentrations of different corrinoids and calculated the corrinoid concentration that resulted in half-maximal growth (EC_50_); the corrinoid with the lowest EC_50_ is considered the most preferred (Fig. 5A and Fig. S4). The corrinoid used for isolation was not the most preferred corrinoid in many cases, likely because the added corrinoid was in 100- to 1,000-fold excess in the isolation medium (Fig. 5B). Further, despite our previous finding that most corrinoids other than B12 have not been detected in this soil (30), we found that all of the isolates can use at least one corrinoid in addition to B12, with B12 preferred by almost all isolates. ADE, CRE and CBI could not be used at any of the tested concentrations by some isolates. Notably, these were the same corrinoids in which we recovered the lowest numbers of isolates. This suggests only some isolates can use the complete corrinoids ADE and CRE as cofactors or salvage CBI to make a complete corrinoid (59–61).

**Figure 5.**
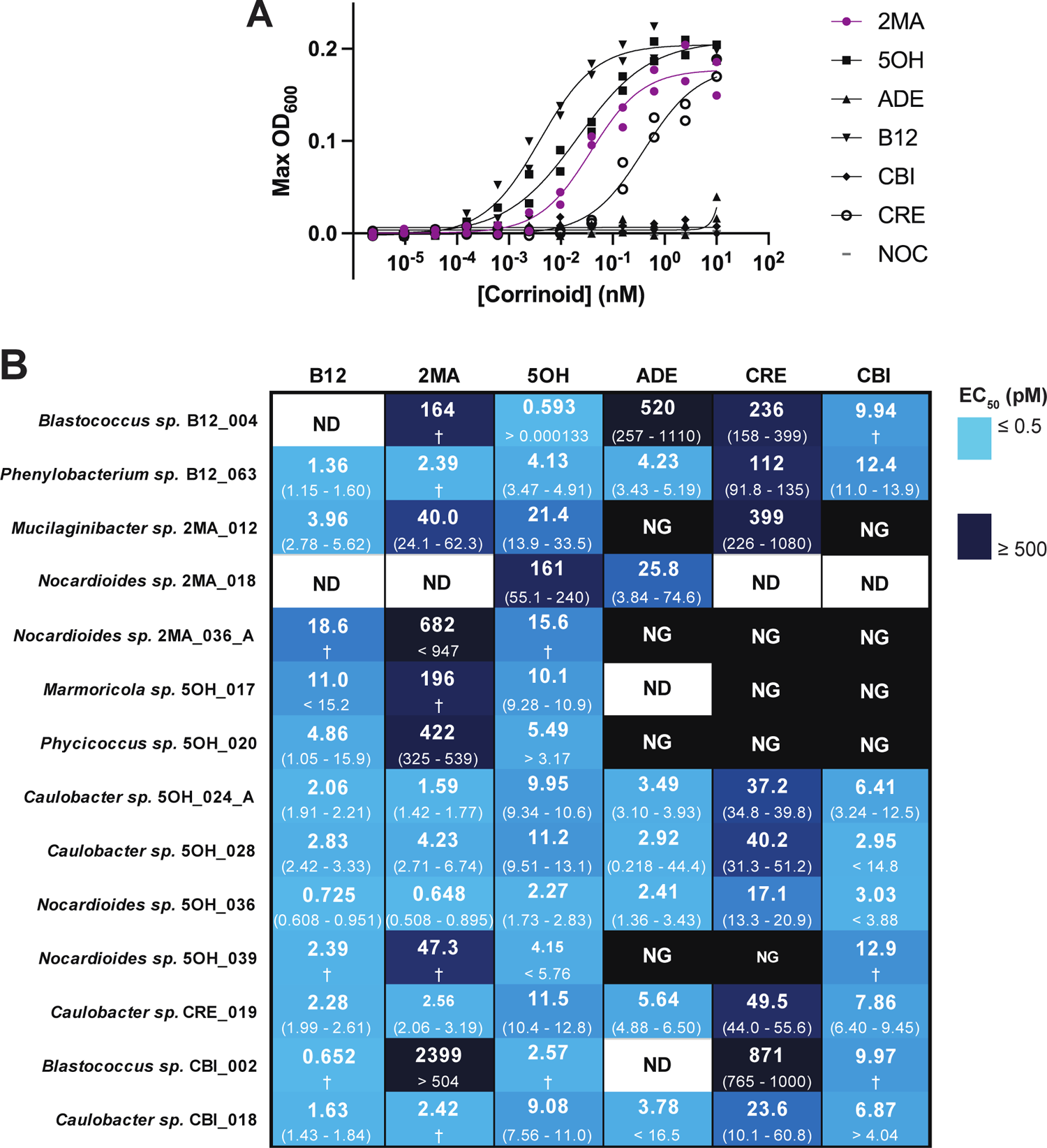
Corrinoid dependence in the isolate collection. (A) Representative dose-response curve belonging to isolate 2MA_012. The corrinoid in which the isolate was recovered is shown in purple. Lines show non-linear fit for each corrinoid. (B) The corrinoid concentrations resulting in half-maximal growth (EC_50_) are shown for all 14 corrinoid-dependent isolates on the six corrinoids used in this study. For each isolate and corrinoid combination, the top number is the EC_50_ and the numbers in parentheses represent the 95% confidence interval as calculated by a four-parameter non-linear fit on GraphPad Prism (v9.5.1). Greater than and less than symbols were used when the upper or lower bound of the confidence interval could not be determined, respectively. NG: No Growth, ND: No Data, † 95% confidence interval not determined. Curve fit data are summarized in supplemental table 7.

### B12 is the main corrinoid produced by isolates in the collection and it is only provided by a subset of producers

Given that dependents need to obtain corrinoids from producers in their community, we sought to determine whether there is compatibility between the corrinoids produced and required by isolates in our collection. To that end, we extracted corrinoids from cultures of 12 fast-growing producers and analyzed them by HPLC. We detected a corrinoid by HPLC in 11 of these producers. The other isolate showed no corrinoid signal when re-tested with the *E. coli* bioassay and was reclassified as inconclusive. Comparison with authentic standards revealed that B12 was the dominant corrinoid synthesized by the 11 tested producers (Fig. 6A).

**Figure 6.**
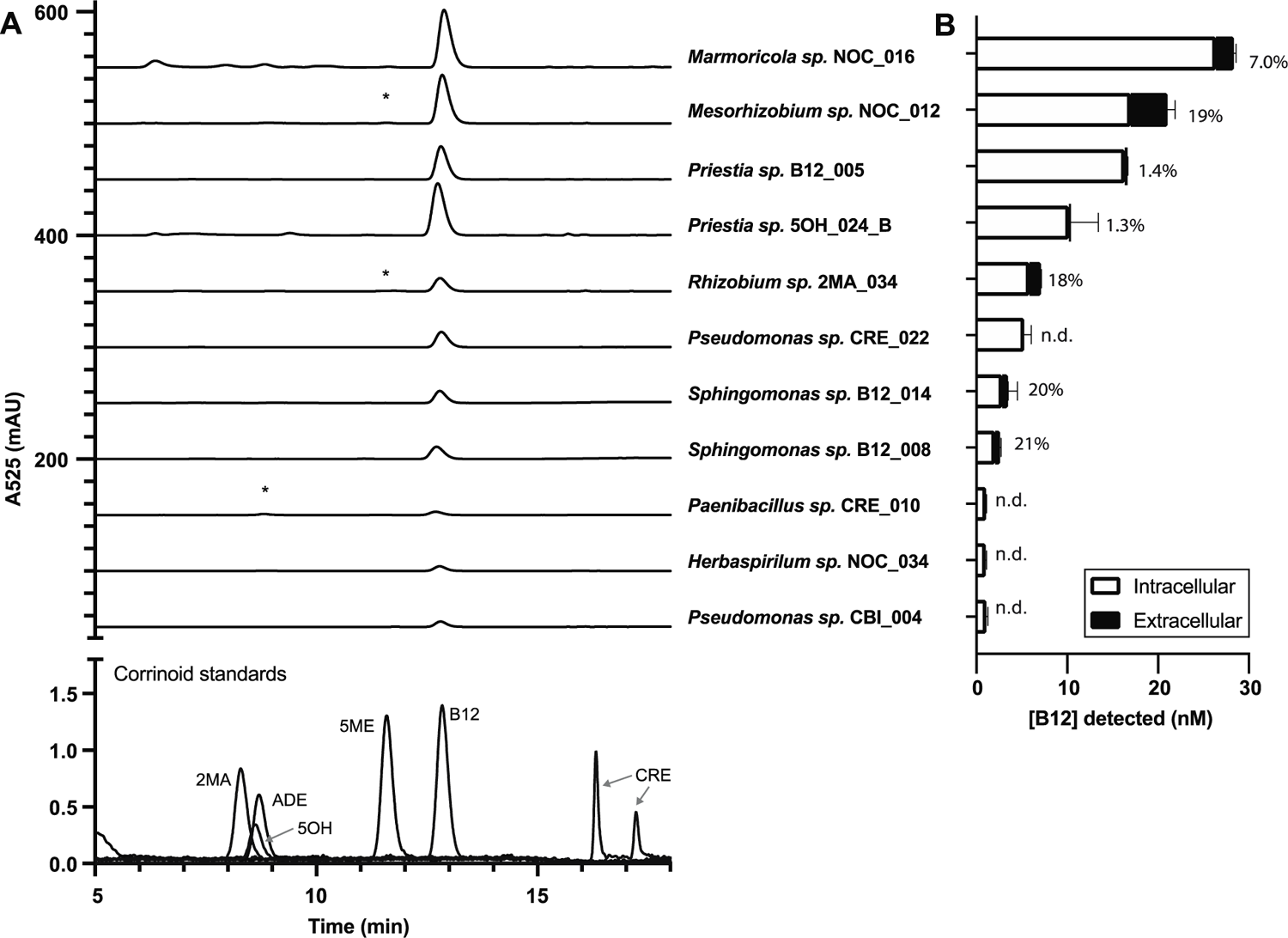
Corrinoid production and providing in the isolate collection. (A) HPLC analysis of corrinoid extracts of 11 selected producers shows B12 is the major corrinoid produced. Authentic corrinoid standards are shown at the bottom. Asterisks denote small peaks that indicate the presence of a second complete corrinoid. (B) Quantification of corrinoids in the cell pellet (intracellular) and supernatant (extracellular) fractions of each isolate as detected by an *E. coli*-based corrinoid bioassay. The percent of corrinoid provided (extracellular corrinoid as a fraction of the total corrinoid) is given to the right of each bar. Bars and error bars show the average and standard deviation of three technical replicates, respectively. n.d., extracellular corrinoid was not detected.

Although it is unknown to what extent dependents acquire corrinoids directly from producer cells, a recent report showed that some producers cultured in the laboratory can release corrinoids into the growth medium (23). We used the *E. coli*-based bioassay to quantify the corrinoids in culture supernatants and cell pellets of the 11 producer isolates. Seven of the producers were found to be “providers” (23) – producers for which corrinoids were present in the culture supernatant – while corrinoids were detected exclusively in the cell pellet fraction in the remaining four producers (Fig. 6B). The amount of provided corrinoid ranged from 1.3 to 21% of the total corrinoid, and the provided amount did not correlate with the total amount of corrinoid produced. The concentrations of provided corrinoid are 1 to 1,000 times higher than the EC_50_ values calculated for the dependents (Fig. 5B, Supplemental Table 6), suggesting that these isolates have the capacity to provide sufficient or excess corrinoid to all of the dependents in our collection. Thus, our measurements of corrinoid production and providing, in the artificial conditions of laboratory culture, coupled with prior genomic studies (25,26), support our hypothesis that corrinoid sharing can occur within the communities of this soil.

## DISCUSSION

Microbial nutritional interactions play pivotal roles in establishing community structure and function. Characterizing and predicting the ecological roles of microbes as nutrient producers and dependents can contribute to the understanding of microbial interaction networks and their influence on the whole community. Here, we investigated the ecological roles of microbes by overlaying experimental and computational approaches. We were able to characterize specific functional roles of bacteria by focusing on a single class of model nutrients, corrinoids, the sharing of which is thought to be widespread in microbial communities (17,25,27,28). The importance of corrinoids for soil bacteria has long been recognized (57,62–64). Here, we report the first systematic isolation and characterization of soil bacteria on corrinoids other than B12, allowing us to consider the ecological roles of corrinoid producers and dependents in the context of soil microbial ecology.

Microbes typically have preferences for different corrinoids that are reflected in their EC_50_ values (17,29,65). These preferences result from corrinoid transport efficiency and the affinity of corrinoids for the enzymes that use them, and what corrinoid dependent processes are in use (65–67). Corrinoid preferences, combined with the availability of corrinoids in a given environment and competition for corrinoids in the community, can impact microbial fitness (17). After experimentally determining the preferences of the corrinoid-dependent bacteria in our collection, we found that our isolates have considerably lower EC_50_ values for their preferred corrinoids than bacteria for which EC_50_ measurements have previously been reported, indicating lower corrinoid concentrations are required for growth (35,48,66,68). The EC_50_ values of our isolates are comparable to those of aquatic algae (29,69), some of which live in environments with corrinoid concentrations in the picomolar range (70) (Fig. 5B and Table S1). The ability to use corrinoids at low concentrations could be a useful adaptation to the soil environment where corrinoids may be limiting due to the physical heterogeneity of soil microbial communities, long distances between cells, and fluctuations in water content throughout the year, making nutrient availability highly variable (13,71,72).

The concentration of corrinoid chosen for the isolation media was four-fold higher than the highest EC_50_ and over 16,000-fold higher than the lowest EC_50_ we measured, which explains why isolates were often recovered in their non-preferred corrinoid and why we detected no taxonomic trends in the corrinoid used for isolation. Despite the presence of excess corrinoid in our isolation media, we recovered fewer corrinoid-dependent isolates than expected (26,57), which may be due to corrinoid-dependent bacteria having additional nutrient dependencies not satisfied by our isolation medium or requiring specific partners for their survival (6). Notably, all of the dependent isolates were able to use corrinoids that have not been detected in this soil (30).

When considering how our classifications of corrinoid metabolism fit into the context of soil microbial ecology, we must consider functional diversity (73). A contemporary question regarding microbiome function relates to whether groups of microbes with shared functions are composed of phylogenetically close organisms or unrelated organisms that share similar metabolic capabilities. Traits such as photosynthesis, methanogenesis, maximum growth rates, and response to soil wet-up tend to be strongly correlated with phylogeny, while others, such as use of specific carbon sources, have weak or no phylogenetic signals (74–76). Here, we found some phylogenetic trends in corrinoid traits, but overall, the distribution of these traits is patchy across the phylogenetic tree, suggesting that gene loss has occurred at various evolutionary points, possibly due to the frequent emergence of corrinoid dependence and independence (9), or that horizontal gene transfer (HGT) is important in sustaining corrinoid biosynthesis and use. Indeed, corrinoid uptake genes in human gut Bacteroidetes are commonly found on mobile genetic elements (77), and *Salmonella typhimurium* and *Lactobacillus reuteri* biosynthesis genes are thought to have been acquired by HGT (78,79). The evolutionary history of corrinoids should be explored further to identify which processes have impacted the biosynthesis and use of these cofactors. Importantly, we were able to carry out the crucial step of validating some genus-level predictions using our isolate collection, in which seven of the genera for which we predicted a genotype were represented (Fig. 4).

Finally, the characterization of this isolate collection provides key insights about corrinoid-based microbial interactions in soil. We found that dependent isolates were all able to use B12, with most preferring it, and the producers we characterized all synthesize B12, indicating compatibility between corrinoid production and preferences of the dependents. Among the producers, however, only some release corrinoid into culture supernatants, suggesting that the corrinoid provider role is fulfilled by a distinct subset of producers (23). Based on our observation that these providers release corrinoids at levels sufficient to support many dependents in laboratory cultures, we speculate that a small fraction of the community disproportionately provides corrinoids to the dependents. In a similar vein, we previously found that amino acid auxotrophs can be supported by producers at a ratio of over 40:1 (80). Because corrinoid release cannot be predicted from genomes, it is necessary to combine genotypic predictions of corrinoid production with phenotypic characterizations when studying interactions. This collection of isolates assembled from the same study site will enable further investigation of corrinoid-based interactions via culture-based studies. For example, the mechanisms of corrinoid release, partner specificity in interactions, and competition for corrinoids among dependent isolates can be explored. Focusing on corrinoids simplifies community interactions to only one nutrient and does not take into account other possible interactions that are prevalent in the soil environment, including those involving other shared nutrients (81,82) or cross-domain interactions (56), which may be affected by environmental fluctuations in the native environment (71). Nonetheless, focusing on this important class of shared nutrients enabled us to study the diversity of metabolic capabilities that may be prototypical of interactions among soil bacteria and provides a framework to expand the study of other nutrient-sharing interactions.

## ACKNOWLEDGEMENTS

We thank Janani Hariharan, Mary Firestone, Britt Koskella, Will Ludington, Victor Reyes-Umana, and all members of the Taga Lab for helpful discussions. We are grateful to Victoria Innocent for providing corrinoids, Heejung Cho and Shi Wang for providing strains, Mary Firestone for allowing us to access her study site at Hopland Research and Extension Center, Katerina Estera Molina for soil sampling support, and Darryl Balderas for computational support. We thank Rebecca Procknow, Dennis Suazo, Janani Hariharan, and Eleanor Wang for critical reading of the manuscript. Sequencing was performed by QB3 Genomics, UC Berkeley, Berkeley, CA, RRID:SCR_022170 and the UC Berkeley DNA Sequencing Facility. This work was supported by the U.S. Department of Energy (DOE), Office of Biological and Environmental Research (BER), Genomic Sciences Program (GSP) Grant DE-SC0020155 (M.E.T.), the GEM Foundation (Z.I.A.A.), the Kase-Tsujimoto Foundation (Z.I.A.A.), and the Sponsored Projects for Undergraduate Researchers program at UC Berkeley (M.A.G.). E.L.B. was supported in part by the DOE, BER, GSP LLNL ‘Microbes Persist’ Soil Microbiome Scientific Focus Area SCW1632. We acknowledge that this work was conducted on the ancestral and unceded land of the Ohlone and Pomo people.

## Data Availability Statement

The sequencing data generated and analyzed during the current study are available in the NCBI GenBank repository under accession numbers OR878823-OR878983. Code generated during the current study is available in https://github.com/zoilaalvarez/corrinoid-metabolism-analysis.

