## Supplemental Materials for "Phylogenetic distribution and experimental characterization of corrinoid production and dependence in soil bacterial isolates"

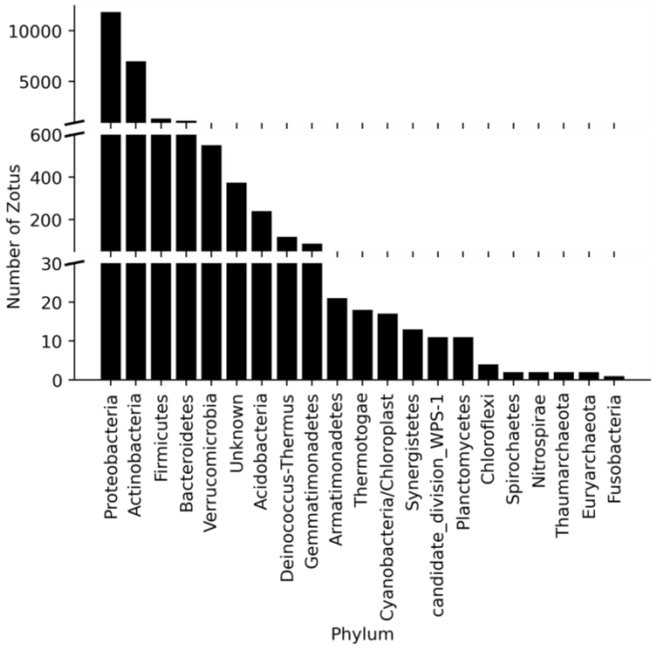


**Figure S1. Phylum classification of microbes cultured by the limiting dilution method.** Data are shown from Illumina sequencing of amplicons of the 16S V4-V5 region in the second isolation batch. 5.8% of these zOTUs were found to be in clonal wells, while the majority of wells contained two or more zOTUs.

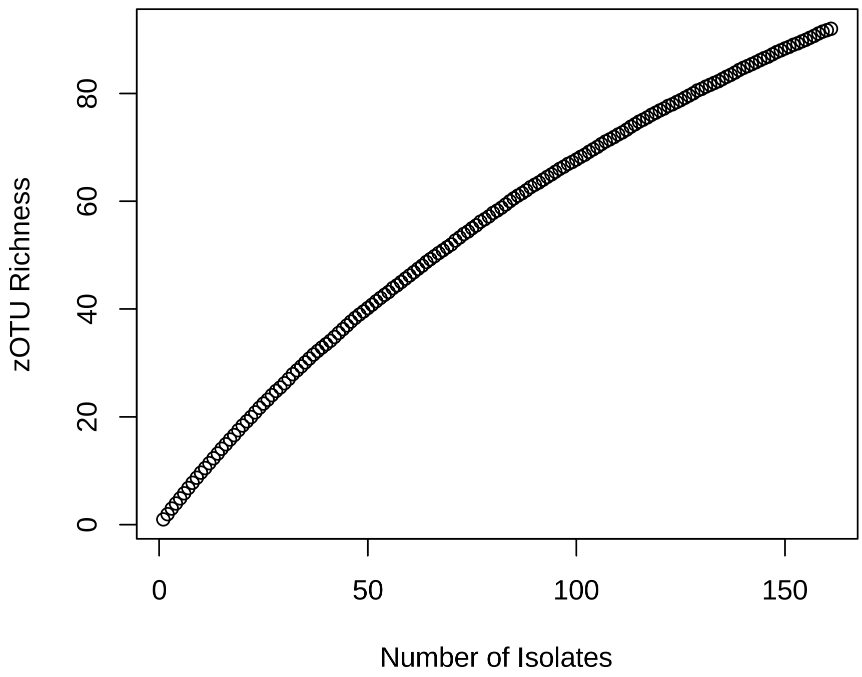


**Figure S2.** The collector's curve for the collection of 161 isolates shows that our isolation effort was far from saturating the bacterial diversity of this soil.

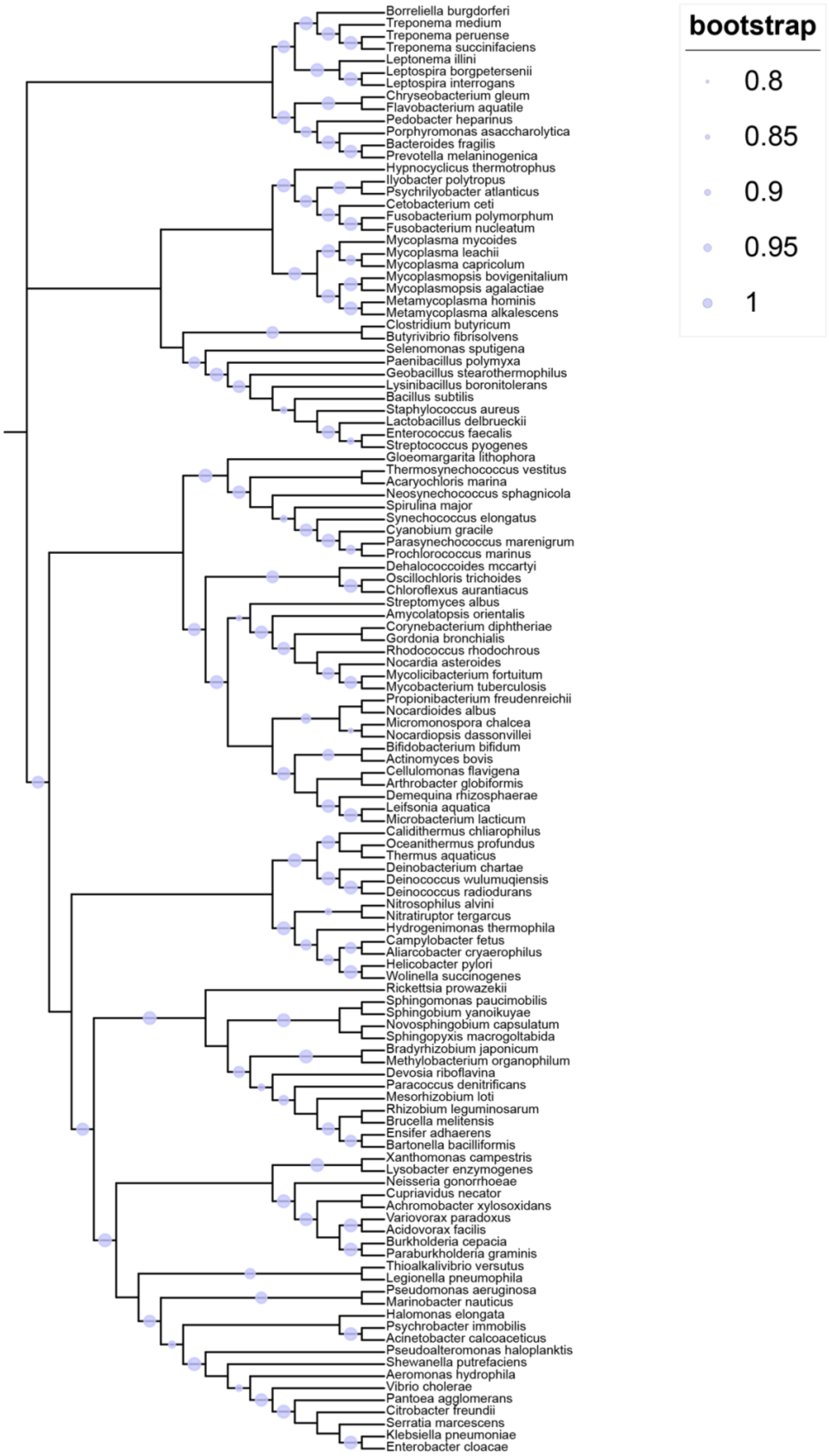


**Figure S3.** Phylogenetic tree of all type species used to generate the pruned tree in figure 4.

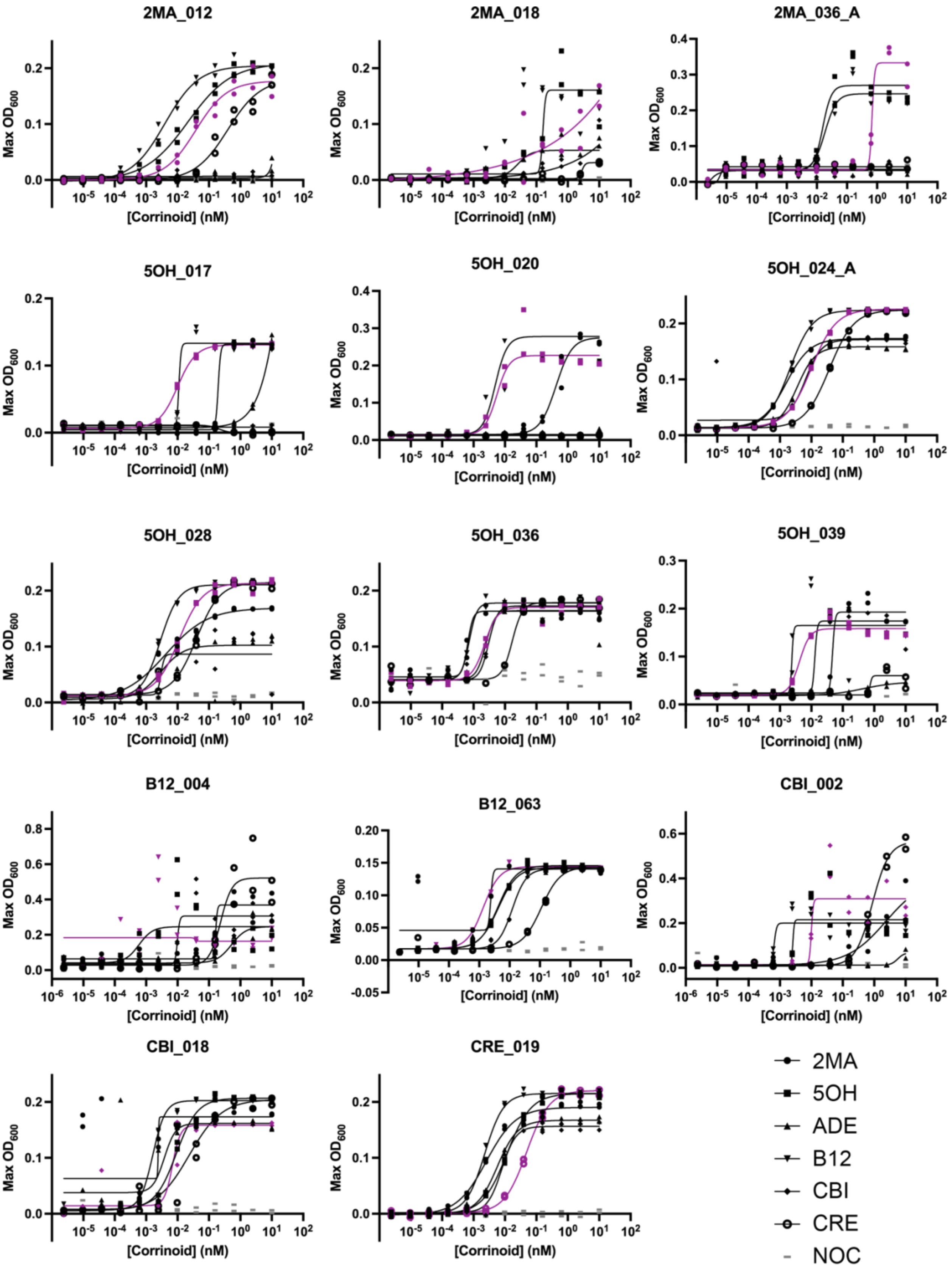
2MA_012
2MA_018
2MA_036_A
5OH_017
5OH_020
5OH_024_A
5OH_028
5OH_036
5OH_039
B12_004
B12_063
CBI_002
CBI_018
CRE_019
Max OD600
[Corrinoid] (nM)
2MA
5OH
ADE
B12
CBI
CRE
NOC

**Figure S4. Corrinoid dose-response curves for all tested dependents reveal widespread preferences for B12.** The corrinoid used for isolation is shown in purple, and the no corrinoid condition is shown in gray. $EC_{50}$ values calculated from these curves are shown in Figure 5B.

**Table S1.** Reported $EC_{50}$ values for B12 in bacteria and aquatic eukaryotic algae.

| **Domain** | **Organism** | **$EC_{50}$** | **Reference** |
|---|---|---|---|
| Bacteria | *Akkermansia muciniphila* | 56.1 pM | (48) |
| Bacteria | *Escherichia coli* Δ*metE* | ~0.1 nM | (35) |
| Bacteria | *Bacteroides thetaiotaomicron* | < 0.4 nM | (66) |
| Bacteria | *Clostridium difficile* | ~1 nM | (68) |
| Eukarya | B12 dependent mutant of *Chlamydomonas reinhardtii* | 28 pM | (29) |
| Eukarya | *Karenia mikimotoi* | 13.1 pM | Tang 2010 PNAS |
| Eukarya | *Aureococcus anophagefferens* | 3.49 pM | (69) |
| Eukarya | *Rhodomonas salina* | 0.36 pM | (69) |
| Eukarya | *Fibrocapsa japonica* | 0.28 pM | (69) |
| Eukarya | *Chattonella marina* | 0.19 pM | (69) |
| Eukarya | *Prorocentrum minimum* | 0.02 pM | (69) |
| Eukarya | *Pavlova lutheri* | ~ 18 pM | (29) |
| Eukarya | *Ostreococcus tauri* | < 70 pM | (29) |
| Eukarya | *Amphidinium carterae* | < 70 pM | (29) |
| Eukarya | *Thalassiosira pseudonana* | < 70 pM | (29) |
| Eukarya | *Aureococcus anophagefferens* | < 70 pM | (29) |
| Eukarya | *Lobomonas rostrata* | < 70 pM | (29) |
| Eukarya | *Euglena gracilis* | < 70 pM | (29) |
